# Up-regulation of bundle sheath electron transport capacity under limiting light in C_4_ *Setaria viridis*

**DOI:** 10.1101/2021.02.10.430537

**Authors:** Maria Ermakova, Chandra Bellasio, Duncan Fitzpatrick, Robert T. Furbank, Fikret Mamedov, Susanne von Caemmerer

## Abstract

C_4_ photosynthesis is a biochemical pathway that operates across mesophyll and bundle sheath (BS) cells to increase CO_2_ concentration at the site of CO_2_ fixation. C_4_ plants benefit from high irradiance but their efficiency decreases under shade causing a loss of productivity in crop canopies. We investigated shade acclimation responses of a model NADP-ME monocot *Setaria viridis* focussing on cell-specific electron transport capacity. Plants grown under low light (LL) maintained CO_2_ assimilation rates similar to high light plants but had an increased chlorophyll and light-harvesting-protein content, predominantly in BS cells. Photosystem II (PSII) protein abundance, oxygen-evolving activity and the PSII/PSI ratio all increased in LL BS cells indicating a higher capacity for linear electron flow. PSI, ATP synthase, Cytochrome *b_6_f* and the chloroplastic NAD(P) dehydrogenase complex, which constitute the BS cyclic electron flow machinery, were all upregulated in LL plants. A decline in PEP carboxylase activity in mesophyll cells and a consequent shortage of reducing power in BS chloroplasts was associated with the more oxidised redox state of the plastoquinone pool in LL plants and the formation of PSII - light-harvesting complex II supercomplexes with an increased oxygen evolution rate. Our results provide evidence of a redox regulation of the supramolecular composition of Photosystem II in BS cells in response to shading. This newly identified link contributes to understanding the regulation of PSII activity in C_4_ plants and will support strategies for crop improvement including the engineering of C_4_ photosynthesis into C_3_ plants.

**Significance statement:** The efficiency of C_4_ photosynthesis decreases under low irradiance causing a loss of productivity in crop canopies. We investigate shade acclimation of a model NADP-ME monocot, analysing cell-specific protein expression and electron transport capacity. We propose a regulatory pathway controlling abundance and activity of Photosystem II in bundle sheath cells in response to irradiance.

## Introduction

C_4_ plants operate a biochemical CO_2_ concentrating pathway that reduces photorespiration by providing higher CO_2_ partial pressure at the site of ribulose bisphosphate carboxylase oxygenase (Rubisco) (Sage and Monson, 1999, von Caemmerer and Furbank, 2003). C_4_ leaves are typically organised in two concentric cylinders around the veins. The vasculature is surrounded by bundle sheath (BS) cells from which radiate mesophyll (M) cells, intertwined by intercellular airspace (Dengler and Nelson, 1999, Lundgren *et al*., 2014). In the cytosol of M cells, CO_2_ is first hydrated to HCO_3_^-^ by carbonic anhydrase (CA) and then fixed by phosphoeno/pyruvate (PEP) carboxylase (PEPC) into C_4_ acids. These diffuse to BS cells where they are decarboxylated either by NADP-ME (NADP-dependent malic enzyme), NAD-ME (NAD-dependent malic enzyme), PEPCK (PEP carboxykinase), or a combination thereof (Furbank, 2011). Many important C_4_ crops, likeZeo *mays, Sorghum bicolor, Saccharum officinarum* and *Setaria italica*, principally produce malate in M cells and use NADP-ME as the predominant decarboxylase. The pyruvate produced by NADP-ME diffuses back to M cells where it is regenerated to PEP by pyruvate orthophosphate dikinase using 2 ATP molecules. For each carboxylation event Rubisco produces two molecules of 3-phosphoglyceric acid which can be exchanged for glyceraldehyde 3- phosphate between BS and M cells, in the so-called triose shuttle (Furbank, 2011). The regeneration of glyceraldehyde 3-phosphate to ribulose 1,5-bisphosphate, the substrate of Rubisco, is exclusively located in BS cells and a key enzyme for the regulation of the regeneration phase is sedoheptulose-bisphosphatase (SBPase) (Harrison *et al*., 2001). At least a half of reducing power requirements of BS cells is met by the decarboxylation of malate, supplying BS cells with NADPH derived from the M, while the ATP required for RuBP regeneration is supplied by photophosphorylation in BS chloroplasts (Bellasio and Lundgren, 2016, Munekage and Taniguchi, 2016).

The partitioning of energy demand between the two cell types is flexible to some extent, due to partial overlapping of the biochemical functionality of M and BS cells, the variable engagement of the decarboxylating pathways and the triose shuttle (Weber and von Caemmerer, 2010, Pick *et al*., 2011, Bellasio and Griffiths, 2014). The energy required in M and BS cells is supplied by two distinct populations of chloroplasts, which harvest light through electron transport chains attuned to the specific biochemical requirements. Photosynthetic electron transport pathways in M are very similar to those in C_3_ chloroplasts (Fig. 1). BS chloroplasts of most NADP-ME plants do not produce NADPH by linear electron flow (LEF), and have little or no Photosystem II (PSII) in their mostly agranal chloroplasts (Chapman *et al*., 1980, Romanowska *et al*., 2008, Furbank, 2011), but are specialised to synthesise ATP, driven by the proton motive force (*pmf*) built-up from cyclic electron flow (CEF) around Photosystem I (PSI) (Nakamura *et al*., 2013, Munekage, 2016). Two CEF pathways operate in BS cells (Fig. 1): the first via the chloroplastic NAD(P) dehydrogenase (NDH) complex and the second via the PROTON GRADIENT REGULATION 5 protein (PGR5) (Munekage *et al*., 2010). NDH complex oxidises ferredoxin and reduces plastoquinone (PQ) while also contributing to establishing *pmf*(Peng *et al*., 2011, Shikanai, 2016, Pan *et al*., 2020). In BS cells of Z. *mays*, the NDH pathway is predominant, whilst PGR5 is more abundant in M cells (Takabayashi *et al*., 2005).

**Fig. 1.**
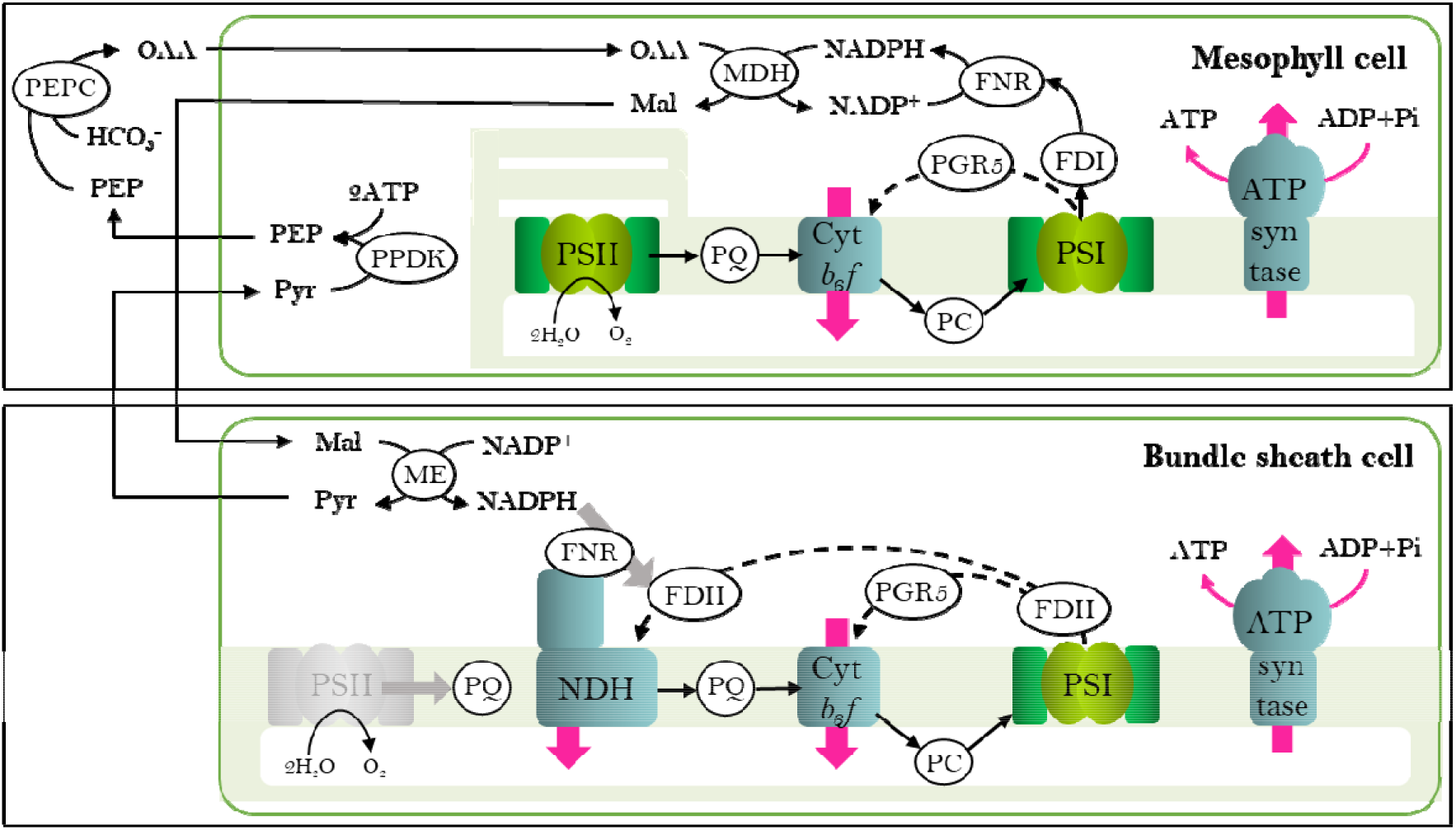
Schematic representation of thylakoid protein complexes and electron transport pathways in mesophyll and bundle sheath chloroplasts of NADP-ME subtype C_4_ plants. PSII, Photosystem II; PQ, plastoquinone; Cyt*b_6_f*, Cytochrome *b_6_f*; PC, plastocyanin; PSI, Photosystem I; NDH, chloroplastic NAD(P) dehydrogenase complex; PGR5, PROTON GRADIENT REGULATION 5; FNR, ferredoxin:NADPH oxidoreductase; FDI, ferredoxin iso-protein I; FDII, ferredoxin iso-protein II; OAA, oxaloacetate; Mal, malate; Pyr, pyruvate; PEP, phosphoeno/pyruvate; PEPC, PEP carboxylase; MDH, malate dehydrogenase; ME, NADP-dependent malic enzyme; PPDK, pyruvate orthophosphate dikinase. Dashed arrows indicate cyclic electron flow pathways, pink arrows indicate proton transfer across the thylakoid membrane and grey arrows indicate potential ways to replenish cyclic electron flow in the bundle sheath chloroplast.

In crop canopies, leaves generally grow in full sunlight and get progressively shaded by new leaves emerging at the top, and up to 50% of net CO_2_ uptake may be fixed by shaded leaves (Baker *et al*., 1988, Long, 1993). C_4_ photosynthesis is known to be sensitive to limiting light intensities as shading decreases the quantum yield for CO_2_ assimilation costing up to 10% of potential carbon gain (Tazoe *et al*., 2008, Pengelly *et al*., 2010, Ubierna *et al*., 2013, Pignon *et al*., 2017). Therefore, long- and short-term acclimations to low light are highly relevant for crop productivity (Bellasio and Griffiths, 2014, Sage, 2014). Reports of PSII activity and grana development in BS cells of NADP-ME plants show great variability depending on the growth conditions and suggest a contribution of electron transport machinery to light acclimation (Andersen *et al*., 1972, Drozak and Romanowska, 2006, Hasan *et al*., 2006, Danila *et al*., 2019). Despite the growing interest in improving C_4_ photosynthesis and attempts to introduce it into C_3_ crop plants (von Caemmerer and Furbank, 2016, Ermakova *et al*., 2019, Ermakova *et al*., 2020b), little is known of environmental responses and regulatory mechanisms of electron transport at the cell-specific level. We investigate the acclimation of thylakoid protein complexes in M and BS cells in a model NADP-ME grass *Setaria viridis* grown under low light (LL), and reveal an up-regulation of the photosynthetic machinery in LL plants, suggesting increased energy requirements, predominantly in BS cells.

## Materials and Methods

### Plant growth conditions

Seeds of *Setaria viridis* (A10 ecotype) were germinated in individual 2 L pots containing garden soil with 2 cm of commercial seed raising mix layered on top (Debco, Tyabb, Australia); both with 1 g L^−1^ of the slow release fertilizer (Osmocote, Scotts, Bella Vista, Australia). Plants were grown in controlled environmental chambers with a 16 h/8 h light/dark period, 28 °C day, 24 °C night, and 60% humidity. Light at 1000 μmol m^−2^ s^−1^ (measured at the pot level) was supplied by halogen incandescent lamps (42W 2800K warm white clear glass 630 lumens, CLA, Brookvale, Australia) and Pentron Hg 4ft fluorescent tubes (54W 4100K cool white, Sylvania, Wilmington, MA). Part of the cabinet was covered with a shade cloth to reduce irradiance to 300 μmol m^−2^ s^−1^. All measurements were performed on the youngest fully expanded leaves sampled before flowering between 15 and 25 days after germination.

### Enzyme activities

Soluble protein was extracted from frozen leaf discs and PEPC activity was measured by spectrophotometric assay after Pengelly *et al*. (2010). The amount of Rubisco active sites was assayed by [^14^C] carboxyarabinitol bisphosphate binding as described in Ruuska *et al*. (2000).

### Western blotting and microscopy

BS strands were isolated following Ghannoum *et al*. (2005). Leaves were homogenised by Omni Mixer (Thermo Fisher Scientific, Tewksbury, MA) during three 10-s cycles at the intensity #7 in 100 mL of ice-cold 50 mM phosphate buffer (pH 7.5) containing 2 mM ethylenediaminetetraacetic acid (EDTA), 5 mM MgCI_2_, 5 mM dithiothreitol and 0.33 M sorbitol. The homogenate was passed through a tea strainer to remove large debris and then BS strands were collected from the filtrate on an 80 μm nylon filter (Merck, Burlington, MA).

For protein isolation frozen leaf discs were ground in ice-cold glass homogenisers and isolated BS strands were ground with a pestle in a chilled mortar, both in 0.5 mL of ice-cold 100 mM trisaminomethane-HCl buffer (pH 7.8) with 25 mM NaCl, 20 mM EDTA, 20 g L^−1^ sodium dodecyl sulphate, 10 mM dithiothreitol and 20 ml L^−1^ protease inhibitor cocktail (Sigma-Aldrich, St Louis, MO). Aliquots were taken for chlorophyll analysis and then extracts were incubated at 65 °C for 10 min and centrifuged at 13,000 *g* for 1 min at 4 °C. Protein extracts were supplemented with 4x XT Sample buffer (BioRad, Hercules, CA) and separated by polyacrylamide gel electrophoresis (Nu-PAGE 4-12% Bis-Tris gel, Invitrogen, Life Technologies Corporation, Carlsbad, CA). Proteins were transferred to a nitrocellulose membrane and probed against photosynthetic proteins with antibodies according to manufacturer’s protocols (Agrisera, Vännäs, Sweden). BS preparations had negligible M contamination, as demonstrated by the immunodetection of PEPC (Fig. S1).

M and BS thylakoids isolation, Blue-Native gel electrophoresis and immunoblotting followed Ermakova *et al*. (2019). Ultrathin sections for transmission electron microscopy (TEM) were prepared after Danila *et al*. (2019) and examined using a Hitachi HA7100 (Hitachi High Technologies, Santa Clara, CA) at 75 kV.

### Chlorophyll, nitrogen and starch

Total chlorophyll was extracted from frozen leaf discs ground using TissueLyser II (Qiagen, Venlo, The Netherlands) or from protein samples in 80% acetone, buffered with 25 mM Hepes-KOH (pH 7.8). Chlorophyll *a* and *b* content was measured at 750 nm, 663.3 nm and 646.6 nm, and calculated according to Porra *et al*. (1989). Chlorophyll *a/b* ratios were determined for BS cells directly from isolated BS strands, while M cell ratios were determined from the mesophyll sap released upon leaf rolling as described in Covshoff *et al*. (2012). The fraction of total leaf chlorophyll in BS cells was calculated as x from: *ax + b* (1-x) = *c*, where *a* is the chlorophyll *a/b* ratio of BS cells, *b* is the chlorophyll *a/b* ratio of M cells, and *c* is the chlorophyll *a/b* ratio of leaf. The fraction of total leaf chlorophyll in BS cells was also calculated based on immunodetection of SBPase (located exclusively in BS cells) from the BS and leaf samples normalised on chlorophyll basis (Fig. S2):

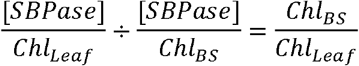

For starch assay, frozen leaf discs were collected 1 h after the light onset and ground with the TissueLyser II, soluble sugars were removed (three extractions with 80% ethanol: incubation for 20 min at 80 °C and centrifuging for 5 min at 13,000 g). Pellets were dried for 15 min at 55 °C and starch was assayed with HK Assay Kit (Megazyme, Bray, Ireland), following the manufacturer’s instructions.

### P700 spectroscopy on bundle sheath strands

Bundle sheath strands were isolated by the differential grinding method (Furbank and Badger, 1983), resuspended in the activity buffer constituted of 10 mM 4-(2-hydroxyethyl)-1- piperazineethanesulfonic acid (Hepes)-KOH (pH 7.4), 2 mM MgCI_2_, 2 mM KH_2_PO_4_, 10 mM KCl, 0.3 M sorbitol to a chlorophyll (*a+b*) concentration of 25-30 μg mL^−1^ and kept on ice. 1 ml of BS suspension was supplied with 10 mM malate, 5 mM dihydroxyacetone phosphate (DHAP), 15 mM ribose-5- phosphate and 100 mM NaHCO_3_ - metabolites required to support CO_2_ assimilation (Furbank and Badger, 1983) - and 200 μM of methyl viologen or 25 μM dichorophenyl-dimethylurea (DCMU) when indicated. Suspension was mixed and BS strands were allowed to sink in a cuvette for 1 min in darkness. After that, the P700^+^ signal was measured from the bottom of the cuvette by Dual-PAM/F (Walz, Effeltrich, Germany). The level of P700^+^ was first recorded in darkness for 1 min and then monitored upon the illumination with red actinic light of 1000 μmol m^−2^ s^−1^. Time constants of P700 oxidation were obtained by exponential fitting in OriginPro 2018b (OriginLab Corp., Northampton, MA).

### Mass spectrometric O2-exchange on bundle sheath strands

Bundle sheath samples (1 ml) were supplied with the metabolites required to support CO_2_ assimilation and filtered via gentle vacuum onto a glass fibre filter (Whatman, Buckinghamshire, UK) through a 10.2 mm aperture. A disc of 10.0 mm was cut from the centre of the filter, saturated with 50 μL of the activity buffer with the metabolites and loaded onto a steel mesh within a 1 mL volume stainless steel cuvette. The cuvette was equilibrated to 25°C via a circulating water bath (Julabo, Seelbach, Germany). The bottom was sealed with a gas permeable Teflon membrane (Hansatech, Norfolk, UK) while the top lid had a quartz window through which halogen light was supplied through a fibre optic and a septum. The cuvette was purged with compressed air and then injected with 2% ^18^O_2_ (99%, Sigma-Aldrich) and 2% CO_2_ (BOC, Sydney, Australia). After 5 min in darkness for equilibration, gross rates of O_2_ production and O_2_ consumption were measured for 5 min in the dark and for 5 min at 1000 μmol m^−2^ s^−1^ via a Delta V Membrane Inlet Mass Spectrometry (MIMS) (Thermo Electron Corp, Bremen, Germany) resolving the evolution of ^16^O_2_ (the product of splitting H_2_^16^O at natural abundance) from consumption of artificially enriched ^18^O_2_. Rates were calculated according to Beckmann *et al*. (2009).

### PSII activity and EPR

PSII activity of thylakoid membranes at a chlorophyll (*a+b*) concentration of 10 μg mL^−1^ and 25 °C, was measured as the rate of oxygen evolution with a Clark electrode (Hansatech Instruments, UK) under white light irradiance (1000 μmol m^−2^ s^−1^) in 20 mM 2-(N-morpholino)ethanesulfonic acid (MES)-NaOH buffer (pH 6.5), 400 mM sucrose, 15 mM NaCl, 5 mM MgCl_2_, with 2 mM ferricyanide and 0.2 mM 2,5-dichloro-1,4-benzoquinone (DCBQ) as electron acceptors.

The flash-induced variable fluorescence decay (Krause and Weis, 1991, Maxwell and Johnson, 2000) was measured to estimate the redox state of the primary quinone electron acceptor in PSII, Q_A_, using a FL3000 dual modulation kinetic fluorometer (Photon Systems Instruments, Drasov, Czech Republic) as described in Volgusheva *et al*. (2016). Samples were measured at a chlorophyll (*a+b*) concentration of 10 μg mL^−1^ and 20 μM DCMU. The fluorescence kinetics were analysed by fitting the multiexponential decay components in Origin 2016 (OriginLab Corp.). Thermoluminescence glow curves, a useful complement to the analysis of the fluorescence kinetics (Vass, 2003, Volgusheva *et al*., 2016), were measured with TL200/PMT thermoluminescence system (Photon Systems Instruments) after Volgusheva *et al*. (2016) at a chlorophyll (*a+b*) concentration of 150 μg mL^−1^ and 40 μM DCMU.

Electron paramagnetic resonance (EPR) of Tyrosine D^•^ from PSII and P700^+^ from PSI (one radical per reaction centre) were quantified with a Bruker EMX-micro spectrometer (Bruker BioSpin, Rheinstetten, Germany) equipped with an EMX-Premium bridge and an ER4119HS resonator using a quartz flat cell as described in Danielsson *et al*. (2004); 15 mM ferricyanide was added when indicated.

### Gas exchange

Net CO_2_ assimilation rate was measured using a portable gas exchange system LI-6800 (LI-COR Biosciences, Lincoln, NE) under red-blue (90%/10%) actinic light. Leaves were equilibrated at 400 ppm CO_2_ in the reference side, flow rate 300 μmol s^−1^, leaf temperature of 28°C, and irradiance of either 1000 or 300 μmol m^−2^ s^−1^.

### Statistical analysis

For all measurements, the relationship between mean values for HL and LL plants was tested using a two-tailed, heteroscedastic Student’s t-test (Microsoft Excel^®^ 2016).

## Results

### Leaf biochemistry

Two weeks-old *S. viridis* grown under low light (LL, 300 μmol m^−2^ s^−1^) developed less tillers and had less leaves than plants grown under high light (HL, 1000 μmol m^−2^ s^−1^, Fig. 2). LL plants had half the dry weight per leaf area and 70% lower starch content compared to HL plants (Table 1). LL leaves contained 63% more chlorophyll (*a+b*) and had higher chlorophyll *a/b* ratio than HL plants (Table 1). At the celltype level, total chlorophyll increased 127% in BS cells and 37% in M cells of LL plants, compared to the corresponding cells in HL plants (Table 1). The chlorophyll *a/b* ratio did not change significantly between M and BS cells; however, BS cells contained 41% of the total leaf chlorophyll in LL plants and 29% in HL plants (Fig. 2).

**Fig. 2.**
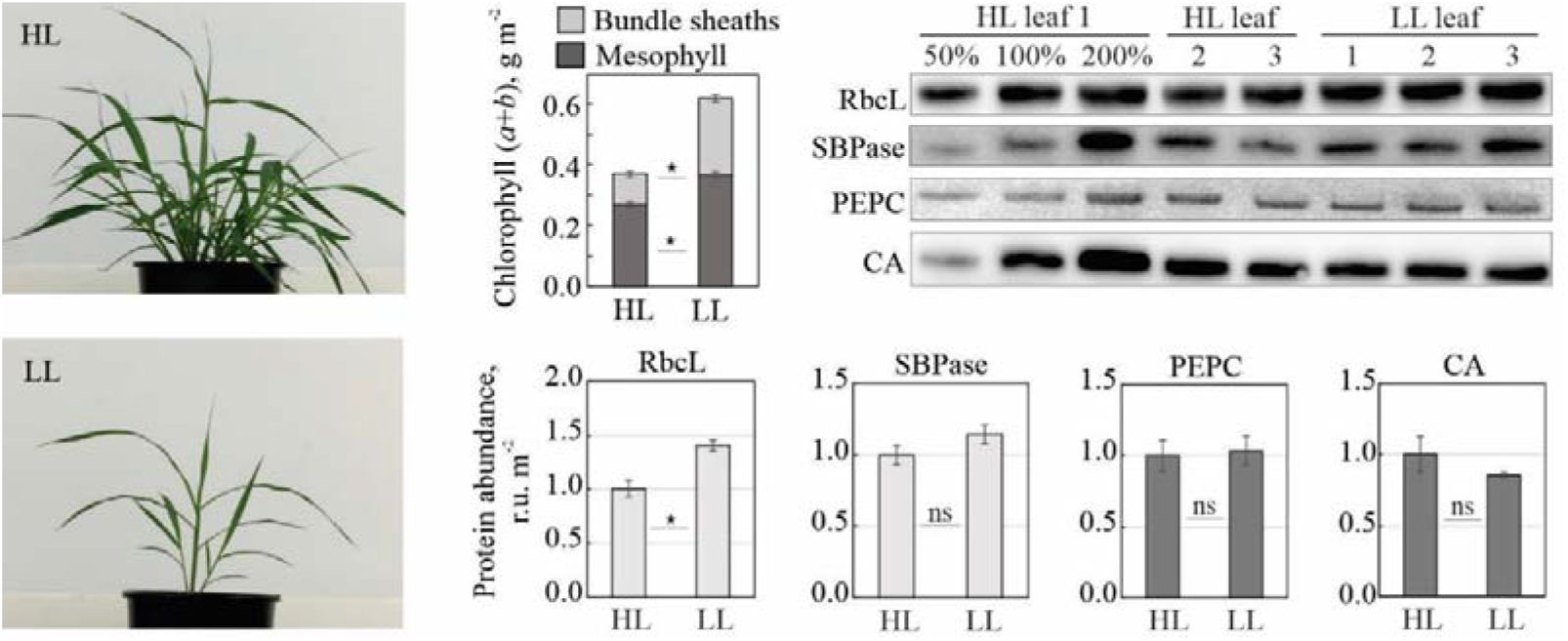
Properties of S. *viridis* grown under high light (HL) and low light (LL). (Left) Plants 16 days after germination. (Top middle histogram) Leaf chlorophyll (*a+b*) distribution between mesophyll and bundle sheaths. (Top right) Immunodetection of large subunit of Rubisco (RbcL), sedoheptulose-bisphosphatase (SBPase), PEP carboxylase (PEPC) and carbonic anhydrase (CA) on leaf area basis. Three biological replicates were loaded for each sample type and a titration series of one of the samples was used for relative quantification. (Bottom right histograms) Relative quantification of protein abundances from the immunoblots. Light grey bars - bundle sheath-specific proteins, dark grey bars - mesophyll-specific proteins. Asterisks indicate statistically significant difference between the two light regimes (P<0.05); mean ± SE; *n =* 3 biological replicates; ns, not significant.

**Table 1.**
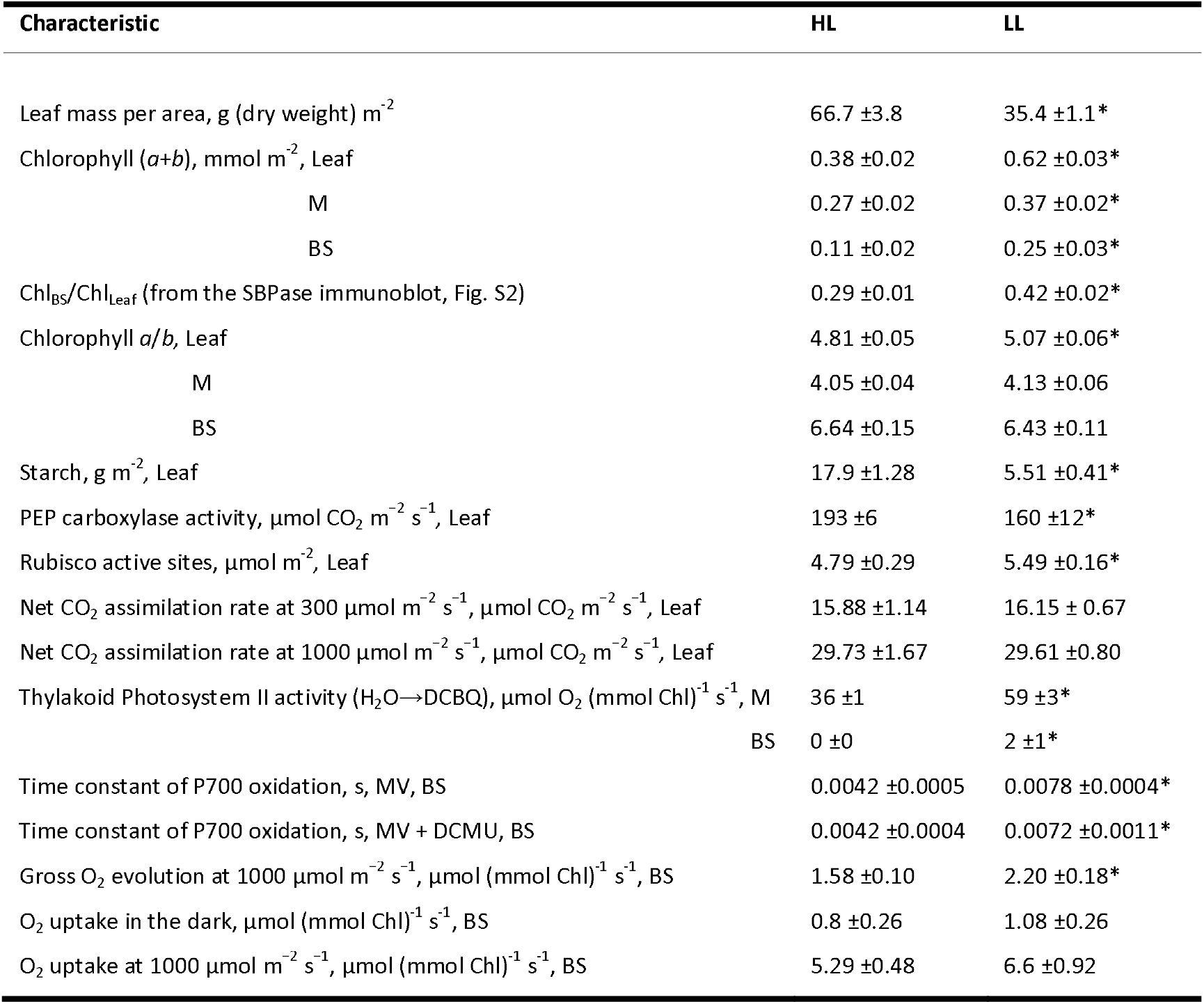
Biochemical and photosynthetic characteristics of leaves, mesophyll (M) and bundle sheaths (BS) cells of S. *viridis* grown at high light (HL) and low light (LL). DCBQ, 2,6-Dichloro-1,4-benzoquinone; MV, methyl viologen; DCMU, dichorophenyl-dimethylurea. Mean ± SE, *n* = 3 biological replicates, except for chlorophyll measurements (n = 6 for M and leaves, *n* = 9 for BS) and BS O_2_ fluxes (n = 5). Asterisks indicate statistically significant difference between the two light regimes (P<0.05).

The abundances of PEPC, CA and SBPase per leaf area did not differ between the two light regimes (Fig. 2), but the activity of PEPC was 20% lower in LL plants (Table 1). The amount of Rubisco active sites was 15% higher in LL plants (Table 1), consistent with the higher abundance of Rubisco large subunit (RbcL) (Fig. 2). Net CO_2_ assimilation did not differ between LL and HL plants (Table 1).

### Cell-level protein abundance

M cells of LL plants had about two-fold more PSI and PSII core subunits (D1 and PsaB, respectively), 1.5-fold more of the Lhcb2 subunit of light-harvesting complex II (LHCII) and 1.3-fold more of the AtpB subunit of ATP synthase than M cells of HL plants (Fig. 3). No difference between HL and LL plants was detected in M abundance of the Rieske FeS subunit of Cyt*b_6_f*, the NdhH subunit of NDH complex, the Lhca1 subunit of light-harvesting complex I (LHCI), the PSII subunit S (PsbS, mediates the fast, ΔpH-regulated component of the non-photochemical quenching, NPQ) and PGR5.

**Fig. 3.**
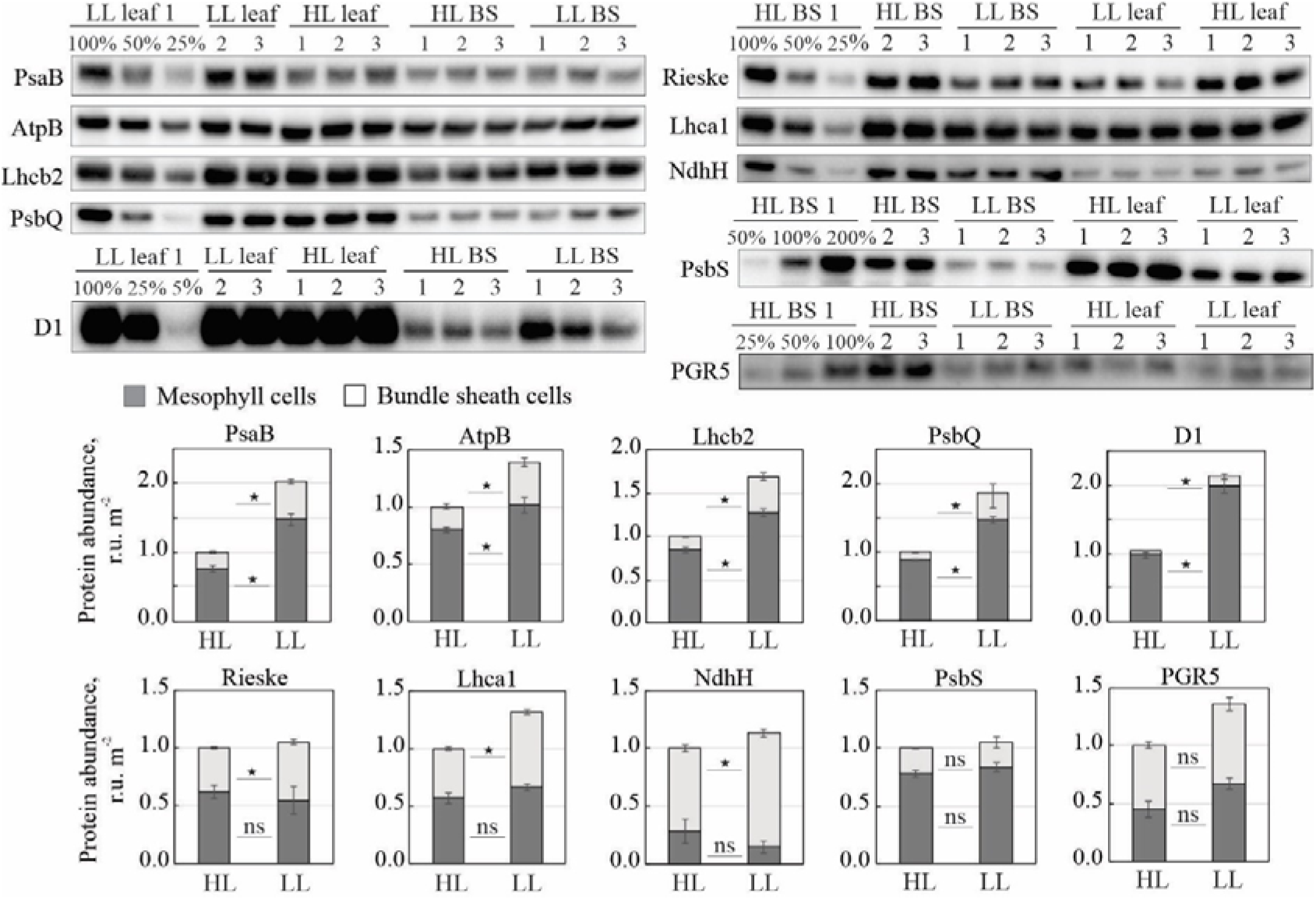
Immunodetection of photosynthetic proteins in *S. viridis* plants grown under high light (HL) and low light (LL). (Top) PsaB (PSI core), AtpB (ATP synthase), Lhcb2 (Light-harvesting complex of PSII), PsbQ (water-oxidising complex of PSII), D1 (PSII core), Rieske FeS (Cytochrome *b_6_f*), NdhH (NDH complex), Lhca1 (Light-harvesting complex of PSI), PsbS (PSII subunit S responsible for photoprotective thermal dissipation) and PGR5 (CEF route) were detected in protein samples isolated from whole leaves (“leaf’) and from bundle sheaths (BS) and loaded on chlorophyll (*a+b*) basis. Three biological replicates were loaded for each sample type and a titration series of one of the samples was used for relative quantification. D1 immunoblot with other dilutions is shown on Fig. S3. (Bottom) Relative quantification of protein abundances in mesophyll (M) and BS cells per leaf area from the immunoblots; HL leaf (M + BS) protein abundances were set to one for each protein. Conversion of relative protein abundances from chlorophyll (*a+b*) to leaf area basis was done using the partitioning of total leaf chlorophyll to BS cells (Table 1). Asterisks indicate statistically significant difference in M or BS cells between the two light regimes (*P* <0.05), mean ± SE, *n =* 3 biological replicates; ns, not significant.

In BS cells, thylakoid protein abundance under LL increased more than in M cells. BS cells of LL plants had about 10-fold more D1, 2.7-fold more PsaB and Lhcb2, 2.3-fold more AtpB, 1.8-fold more Lhca1, 1.7-fold more NdhH, and 1.3-fold more Rieske per leaf area than HL BS cells (Fig. 3). PsbS and PGR5 did not differ significantly between HL and LL BS cells (Fig. 3). BS/M distribution remained similar between HL and LL plants for PsaB, D1, Lhca1, Lhcb2, AtpB, Rieske, PsbS and PGR5, whilst the BS apportioning of NdhH shifted from about 70% in HL plants to about 90% in LL plants. The BS-portion of D1 increased from 3% in HL plants to 8% in LL plants (Fig. 3). For both HL and LL plants, the apportioning of PsbQ, to BS cells was the same or higher than the apportioning of leaf D1 to the BS (Fig. 3).

### PSI/PSII ratio

For M thylakoids of both LL and HL plants, the PSI/PSII ratio was 1.06 (Fig. 4, M LL), similar to C_3_ plants (Danielsson *et al*., 2004). BS thylakoids showed a decreased Tyrosine D^•^ signal (Fig. 4, green spectra) from which a PSI/PSII ratio of 8.0 and 2.6 was calculated for HL and LL respectively.

**Fig. 4.**
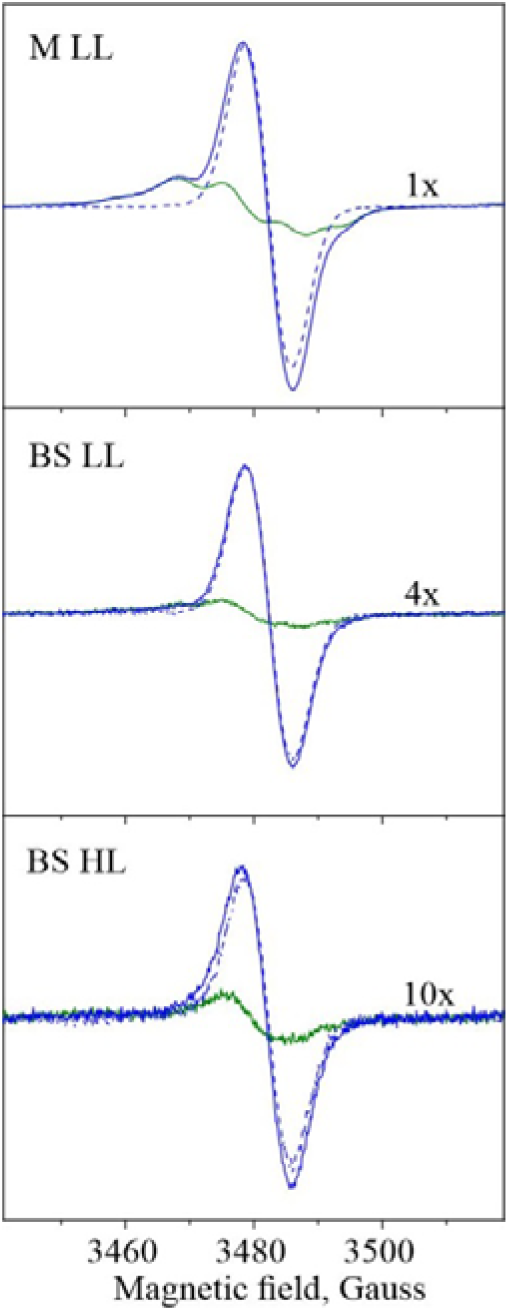
Electron paramagnetic resonance (EPR) quantification of PSI/PSII ratios from mesophyll (M) and bundle sheath (BS) thylakoid membranes of S. *viridis* grown at high light (HL) or low light (LL). Spectra shown are from Tyrosine D radical (green line) oxidised with 15 mM ferricyanide (PSII, blue line) and deconvoluted P700^+^ radical (PSI, dotted blue line). The amplitude of Tyrosine D radical signal in the BS was multiplied for better visibility. EPR measurements were conducted at the microwave frequency 9.76 GHz, microwave power 8 mW, modulation amplitude 5 G and room temperature. The amplitude of Tyrosine D^•^ signal (green line) was multiplied for better visibility. Data shown are from a representative experiment from three independent thylakoid isolations; M HL samples showed similar characteristics to M LL and are not shown.

**Fig. 5.**
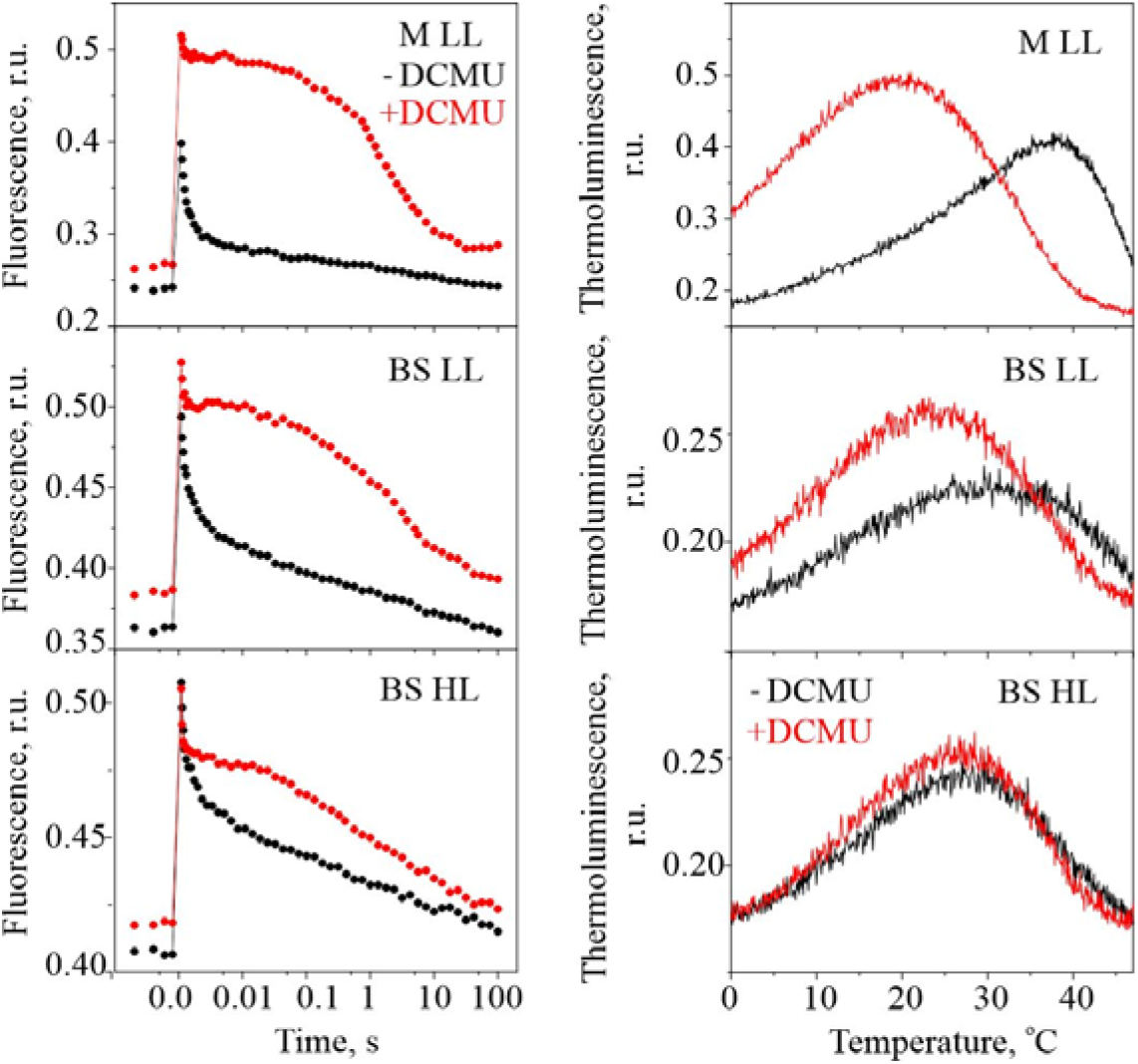
Electron transfer properties of PSII from mesophyll (M) and bundle sheath (BS) thylakoid membranes of S. *viridis* grown at high light (HL) or low light (LL). (Left) Flash-induced fluorescence decay kinetics of thylakoids in the absence (black trace) and in the presence (red trace) of 20 μM DCMU. (Right) Thermoluminescence measurements of thylakoids in the absence (black trace) and in the presence (red trace) of 40 μM DCMU. Data shown are from a representative experiment from three independent thylakoid isolations; M HL samples showed similar characteristics to M LL and are not shown.

**Fig. 6.**
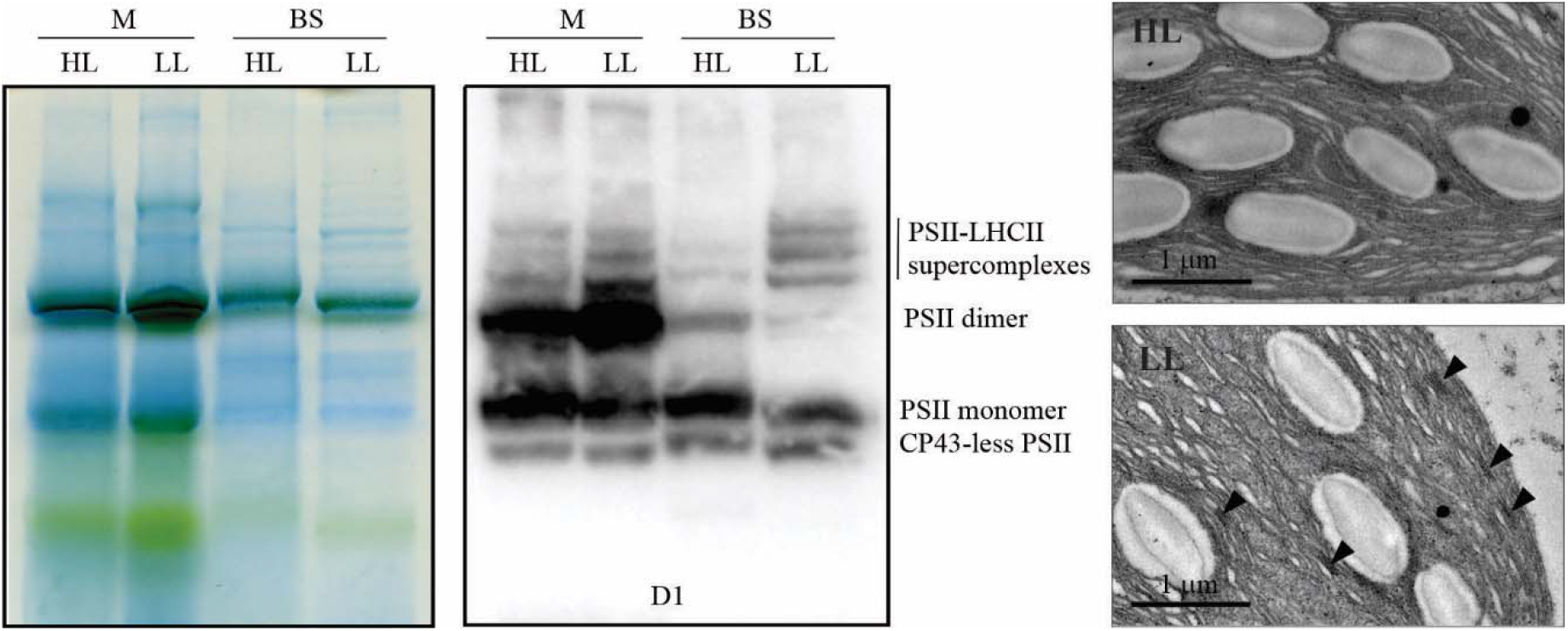
Properties of the thylakoid membranes from S. *viridis* plants grown at high light (HL) or low light (LL). (Left) Blue-Native gel electrophoresis of the protein complexes isolated from mesophyll (M) and bundle sheath (BS) thylakoids and supramolecular composition of PSII analysed by immunodetection of D1; 10 μg of chlorophyll (*a+b*) loaded for each sample. (Right) TEM micrographs of BS chloroplasts, arrows indicate grana formations in LL plants. Scale bar, 1 μm.

### Photosystem II activity

M thylakoids of HL plants had 40% lower O_2_ evolution compared to LL plants (Table 1). Evolution of O_2_ from BS thylakoids of HL plants was below the detection limit and BS thylakoids of LL plants had 11% of the O_2_ evolution of M thylakoids (Table 1).

In the flash-induced variable fluorescence decay, a saturating flash fully reduces Q_A_ resulting in maximal fluorescence yield (Fig. 5, left panels) and the following fluorescence decay kinetics depends on the Q_A_^-^ re-oxidation (Vass *et al*., 1999, Mamedov *et al*., 2000). Three exponential decay components differing by speed were obtained (t_1_, t_2_ and t_3_ in Table S1). The first component represents the reoxidation of Q_A_^-^ by the plastoquinone that is already bound at the Q_B_ site of PSII; the second component represents the re-oxidation by the plastoquinone which has to bind to the Q_B_ site from the PQ pool; the third component reflects a recombination between Q_A_^-^ and the S_2_ state of the water-oxidising complex on the donor side of PSII (Fig. 5, left panels, black traces). In the presence of DCMU, blocking all forward electron transfer from Q_A_^-^ to Q_B_ (Fig. 5, left panels, red traces), the decay phase can be resolved in two slow components (t_2_ and t_3_ in Table S1). M thylakoids from LL and HL plants had similar fluorescence kinetics and therefore only M LL are shown (Fig. 5).

In M thylakoids from LL plants the first component represented 73% of the maximal fluorescence amplitude, the second 14%, and the third 13% (Table S1). With DCMU the first component was 27%, and the second 72%. In BS thylakoids from LL plants, the initial fast fluorescence decay became slower (Fig. 5, left panels, BS LL, black trace) and, as a consequence, the recombination phase nearly doubled in amplitude (21%, Table S1). In HL BS thylakoids this slowing down was even more pronounced (Fig. 5, left panels, BS HL, black trace): the two fast phases were twice and four times slower than in the M thylakoids (Table S1). This slow forward electron transfer kinetics from Q_A_^-^ indicates that the PQ pool was more reduced in BS thylakoids than in M thylakoids, and more so in HL BS thylakoids.

Thermoluminescence measurements also report on the redox state of PSII (Vass, 2003, Volgusheva *et al*., 2016). In samples containing active PSII centres after a single flash, characteristic B-band which reflects recombination between Q_B_^-^ and the S_2_ state of the water-oxidising complex is observed. This is shown for LL M thylakoids where the thermoluminescence band is peaked at 38 □C (Fig. 5, right panels, M LL, black trace). When forward electron transfer from Q_A_^-^ is blocked, for example upon addition of DCMU, different Q-band which reflects Q_A_^-^ → S_2_ state recombination is observed. In M thylakoids this band was observed at 19 □C (Fig. 5, right panels, M LL, red trace). Therefore, these two bands report on two extreme cases where the electron transfer on the acceptor side of PSII is functional or not.

In LL BS thylakoids (Fig. 5, right panels, BS LL) both B- and Q-bands were shifted closer to each other (32 □C and 24 □C, respectively). Furthermore, in HL BS thylakoids both bands merged to show the same peak temperature at 27 □C in the absence or presence of DCMU (Fig. 5, right panels, BS HL). This is another indication of the modification of the redox state of Q_A_ and impaired electron transfer on the acceptor side of PSII (Volgusheva *et al*., 2016), more in HL BS thylakoids. Changes in the redox state of Q_A_ reflect changes in the redox equilibrium Q_A_ ⇆ Q_B_ ⇆ PQ pool and indicate that while in LL BS cells a small pool of Q_B_ was still available to accept electrons from PSII, in HL BS cells no more PSII acceptors were available.

### Supramolecular organisation of PSII and thylakoid structure

In M thylakoids of HL plants, PSII-LHCII supercomplexes and PSII dimers were a lower fraction of all PSII complexes, compared to LL plants. In HL BS thylakoids, PSII monomers were prevalent and PSII-LHCII supercomplexes and dimers were a minor fraction of all PSII complexes (Fig. 6, left panels). BS cells of LL plants had higher proportion of PSII-LHCII supercomplexes and less PSII monomers compared to HL plants. Whilst HL BS thylakoids did not have any grana formations, in BS of LL plants rudimentary grana were observed as regions of appressed thylakoid membranes (Fig. 6, right panels, pointed by arrows).

### Electron transport pathways in isolated bundle sheath cells

To clarify the contribution of PSII and other electron pathways to reduction of PSI in BS cells (Fig. 1), we monitored the kinetics of P700 oxidation in isolated BS strands (Fig. 7). Upon the illumination, after the initial rise, P700^+^ signal quickly returned to the dark level in both LL and HL samples. We attributed this phenomenon to active CEF returning electrons to the PQ pool to reduce P700^+^. When methyl viologen (MV, a PSI acceptor effectively preventing CEF) was added, P700^+^ quickly reached the steady state (Fig. 7) and the time constant of oxidation was almost doubled in LL plants compared to HL plants (Table 1). Application of DCMU in addition to MV did not change the kinetics of P700 oxidation in either of BS samples significantly (Fig. 7, Table 1). However, the time constant of oxidation was still significantly higher in LL plants than in HL plants (Table 1).

**Fig. 7.**
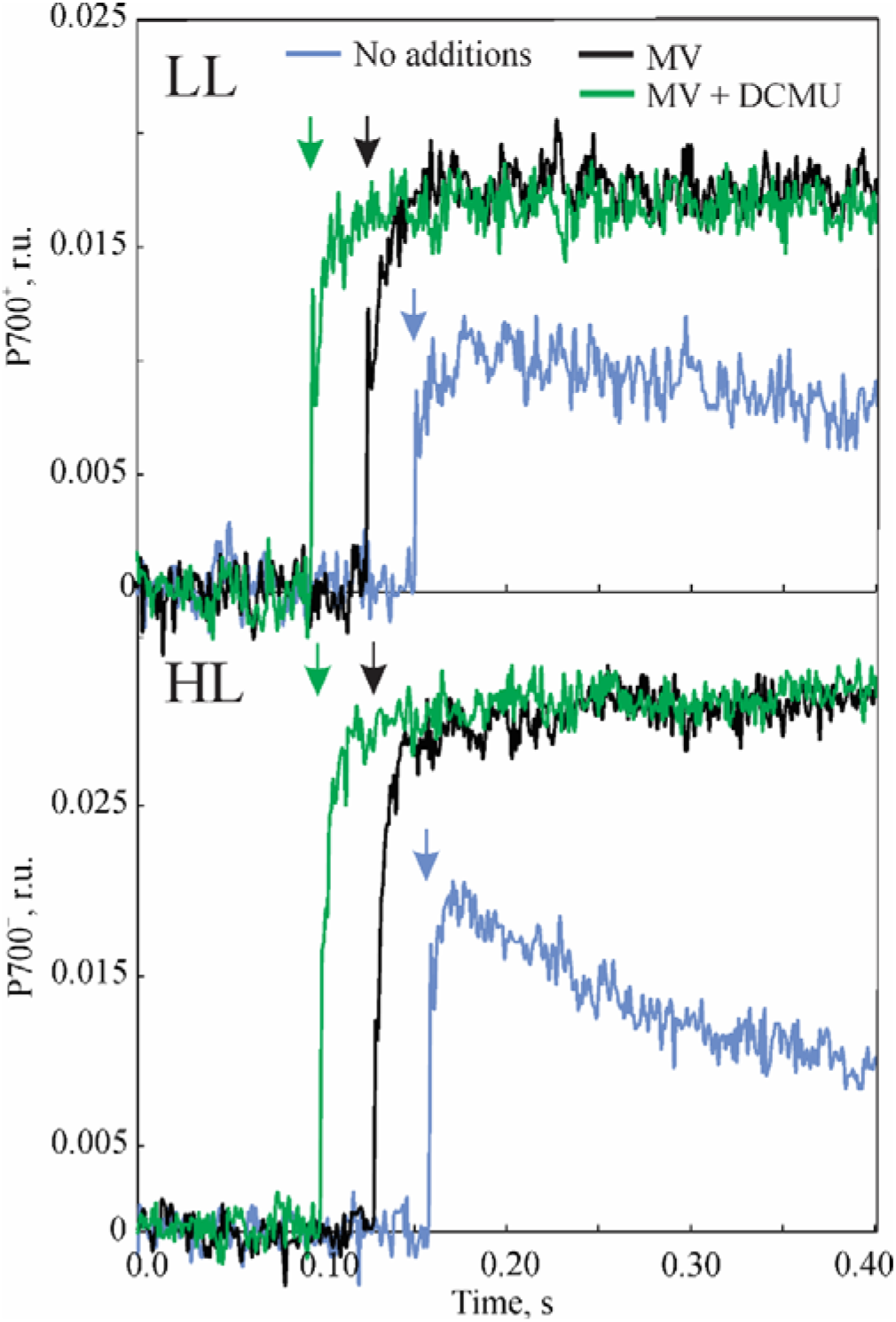
Kinetics of P700 oxidation measured on bundle sheath (BS) strands isolated from S. *viridis* plants grown under high light (HL) and low light (LL), in the absence or presence of 200 μM methyl viologen (MV) and 25 μM DCMU as indicated. All BS preparations were supplied with 10 mM malate, 15 mM ribose-5-phosphate, 100 mM NaHCO_3_ and 5 mM dihydroxyacetone phosphate to support CO_2_ assimilation. Arrows show the beginning of illumination with red actinic light at 1000 μmol m^−2^ s^−1^. Chlorophyll (*a+b*) concentration of BS samples was 25-30 μg mL^−1^. Time constants of exponential fitting of the curves are shown in Table 1.

Isolated BS strands were also assayed for gross O_2_ production and uptake by MIMS at 1000 μmol m^−2^ s^−1^. LL BS cells had gross O_2_ evolution rate of 2.20 μmol (mmol Chl)^−1^ s^−1^, significantly higher than that of HL BS cells of 1.58 μmol (mmol Chl)^−1^ s^−1^ (Table 1). The rates of gross O_2_ uptake in the dark and under irradiance were similar between LL and HL BS cells (Table 1). O_2_ uptake detected in BS cells was not supported by the plastid terminal oxidase (PTOX) as it was only present in M cells (Fig. S4).

## Discussion

We investigated acclimation of the electron transport machinery in *S. viridis*, a model monocot from the Panicoideae subfamily, which affiliates with important NADP-ME crops like *Z. mays, S. bicolor, S. officinarum and S. italica*. LL plants had higher leaf chlorophyll content similar to *Z. mays* (Drozak and Romanowska, 2006, Kromdijk *et al*., 2010) and higher apportioning to the BS (circa 40%) than HL plants (29%), in line with the NADP-ME species *S. bicolor* and *Cencris ciliaris* (Ghannoum *et al*., 2005). This resulted in an increased leaf chlorophyll *a/b* ratio (Fig. 2, Table 1), in contrast to the typical response to shading of C_3_ species upregulating chlorophyll *b*-binding antennae (Boardman, 1977, Sage and McKown, 2006).

BS cells of LL plants upregulated their capacity for LEF, compared to HL plants. The PSI/PSII ratio decreased from 8 (relative to that in the M) in HL plants to 2.6 in LL plants, and the abundance of D1 and PsbQ proteins and PSII activity increased accordingly (Fig. 3, Fig. 4, Table 1). Measured BS PSII activity was higher in LL plants in both isolated BS strands and BS thylakoids (Table 1). Gross O_2_ uptake was detected in HL BS strands through MIMS but not in BS thylakoids trough the polarographic method due to a higher sensitivity of the first. Interestingly, the contribution of PSII to PSI reduction in isolated BS cells was not significant but P700^+^ was oxidised more slowly in LL plants even in the presence of MV and DCMU, blocking CEF and PSII activity (Fig. 7, Table 1). These results are consistent with an existence of another pathway donating electrons to PSI which is more active in LL plants.

A reduction of the PQ pool using stromal reductants mediated by NDH could be responsible for the slower oxidation of P700 in LL BS cells. This is in line with the higher abundance of NDH detected in BS cells of LL plants compared to HL plants (Fig. 3). NDH from Z. *mays* BS copurifies with FNR (Funk *et al*., 1999) and might therefore obtain electrons from stromal NADPH derived from malate decarboxylation via ferredoxin, and, consequently, reduce the PQ pool. In support of this idea, differently from M cells, BS cells of Z. *mays* contain the specific ferredoxin iso-protein FDII and the membrane-bound form of FNR, which has a higher affinity for oxidised ferredoxin and may have evolved to catalyse the reduction of ferredoxin using NADPH, similar to the root FNR (Matsumura *et al*., 1999, Goss and Hanke, 2014).

Increased PSII activity and NDH abundance in LL BS cells could also contribute to upregulation of CEF capacity since the reduction of the PQ pool by LEF and conceivably via NDH, maintains CEF by replenishing electrons leaking to various sinks around PSI (Ivanov *et al*., 2005). Direct CEF measurements in BS cells are complicated by the mutual dependence between M and BS cells and a large proportion of CEF/LET. To overcome experimental limits, we modelled the cell-specific electron transport chains of LL and HL plants and showed that LL BS cells required higher CEF rate than HL plants to sustain the same assimilation rate (see companion paper). The detected changes of protein abundance suggested an increased ATP demand in BS cells of LL plants which is also consistent with an increased CEF capacity. The concurrent upregulation of LHCI and LHCII subunits under LL (Fig. 3) pointed to an increased light-harvesting in the BS. Further, LL plants increased abundances of PSI, Cyt*b_6_f*, NDH and ATP synthase (Fig. 3), which constitute the BS CEF machinery. NDH abundance was previously found to correlate with the ATP requirements of cell types in C_4_ plants (Takabayashi *et al*., 2005); it could contribute to ATP production in BS cells by both cycling electrons around PSI and replenishing the PQ pool with electrons from stromal reductants. Interestingly, PGR5-mediated CEF route was not significantly up-regulated in LL plants and, as suggested by PGR5 overexpression in *Flaveria bidentis* (Tazoe et al., 2020), it is more likely involved in photoprotection of PSI in both M and BS cells than in production of extra ATP in BS cells.

The more oxidised PQ pool in LL BS thylakoids, as revealed by the faster re-oxidation of Q_A_^-^ (Fig. 5), is consistent with the lower intercellular malate flux in LL plants. If NDH mediated the reduction of PSI from stromal reductants, as proposed above, the redox state of the PQ pool would correspond to the stromal availability of NADPH and ultimately to the influx of malate to BS cells. Consistently, we measured in LL plants a lower activity of PEPC (Table 1), which is responsible for the production of C_4_ acids in M cells, and light-activated (Bailey *et al*., 2007). Light regulation of PEPC allows matching the supply of NADPH to BS cells with the BS capacity for ATP production upon changes of irradiance (Pfeffer and Peisker, 1998, von Caemmerer and Furbank, 2003).

The redox state of the PQ pool could affect supramolecular organisation of PSII to facilitate gross O_2_ evolution when the malate supply to BS cells is limited, *i.e*. in LL conditions (Fig. 6). PSII-LHCII supercomplexes, prevalent in LL BS thylakoid, are more stable and capable of higher O_2_ evolution rates than monomers (Hankamer *et al*., 1997, Danielsson *et al*., 2006). It was shown in *Z. mays, S. bicolor* and *F. bidentis* that oxygenic activity of PSII in the BS could be downregulated by reducing the abundance of small subunits PsbP, PsbQ and PsbR required to stabilise the O_2_-evolving complex (Höfer *et al*., 1992, Meierhoff and Westhoff, 1993). However, it was noted by Romanowska *et al*. (2006) that the loss of these subunits may result from the enzymatic treatment of BS strands, while, in our untreated BS preparations, PsbQ was more abundant than D1 both for HL and LL plants, suggesting that PsbQ was not limiting O_2_ evolution (Fig. 3).

Since PSII-LHCII supercomplexes provide a binding site for the grana formation between the stroma-exposed protein residues of two complexes (Albanese *et al*., 2020), the rudimental grana present in LL BS cells (Fig. 6) could have formed because more supercomplexes were available in response to the lower redox state of the PQ pool (Fig. 5). In C_3_ plants, the disassembly of PSII-LHCII supercomplexes - during the LHCII state transitions or for the repair process - is controlled by the STATE TRANSITION 8 (STN8) kinase, phosphorylating the PSII core proteins D1, D2, and CP43 (Tikkanen *et al*., 2008, Dietzel *et al*., 2011). The activity of STN8 requires reduced Cyt*b_6_f* and was proposed to be regulated through the redox state of the PQ pool (Rochaix, 2013, Betterle *et al*., 2015). Although in C_4_ chloroplast proteome studies STN8 was identified with confidence only in M cells (Majeran *et al*., 2008), the phosphorylated D1 was detected in the BS of Z. *mays* (Rogowski *et al*., 2019). Therefore, it is conceivable that an active influx of malate causing overreduction of the PQ pool in HL BS cells could increase STN8 activity and lead to the disassembly of PSII-LHCII supercomplexes and consequently, lower PSII activity. Monomeric PSII with limited oxygenic activity, detected in HL BS thylakoids (Fig. 6, Table 1), was observed in other NADP-ME species (Majeran *et al*., 2008, Hernández-Prieto *et al*., 2019) and might have a role in photoprotection of BS cells allowing the dissipation of excess absorbed light via photoinhibitory quenching (Malnoë, 2018). Alternatively, plants might retain monomeric PSII in BS cells to ensure fast assembly of PSII-LHCII supercomplexes during the progressive shading of leaves in the canopy. Whilst an adjustment of the PSII supramolecular composition could represent a short/medium-term response to irradiance, the persistent overreduction of the PQ pool in HL BS cells may have driven a long-term response of decreasing overall PSI/PSII ratio mediated by regulation of gene expression (Pfannschmidt *et al*., 1999).

The adjustment of PSII activity through the redox state of the PQ pool explains contrasting acclimation strategies of NADP-ME plants, as well as the increased LEF and the granal organisation of BS chloroplasts in NAD-ME plants (Ghannoum *et al*., 2005, Ueno *et al*., 2005). The rate of NADPH supply to the BS can be decreased by diverting a part of the oxaloacetate produced by PEPC to aspartate, which then moves to the bundle sheath, is reconverted to oxaloacetate and decarboxylated through PEPCK or by NADP-ME after re-reduction to malate (Furbank, 2011). The reduction state of the PQ pool may depend on the activity of these redundant decarboxylating pathways, explaining the great variability of PSII content in BS cells of NADP-ME plants (Chapman and Hatch, 1981, Meister *et al*., 1996) and the variability in the acclimation of light reactions in NADP-ME plants (Rogowski *et al*., 2019). The universal nature of PQ-mediated regulation of light reactions gives hope that BS chloroplasts of C_3_ plants engineered with the NADP-ME C_4_ pathway will adjust their PSII activity according to the amount of NADPH provided by malate decarboxylation (Ermakova *et al*., 2020a, Ermakova *et al*., 2020b).

## Conclusion

We examined the effect of growth at low irradiance on thylakoid composition in M and BS chloroplasts of S. *viridis. We* showed that S. *viridis* grown under LL upregulated photosynthetic capacity of BS cells, suggesting an increased energy demand in that compartment, although maintaining strikingly similar CO_2_ assimilation rates to HL plants. Detailed characterisation of PSII activity in BS cells revealed that, similarly to C_3_ plants, the redox state of the PQ pool could affect the composition of the electron transport chain in response to irradiance. In a companion paper, we address a contribution of the detected electron transport changes to leaf-level photosynthetic efficiency, providing fodder for improving yield of C_4_ crops and engineering C_4_ pathway to C_3_ plants.

## Author contributions

ME, SvC, and RTF conceived the project; ME, FM and DF performed experiments; ME, CB and FM analysed data and wrote the article with contribution of SvC and RTF.

## Acknowledgements

We thank Soumi Bala for help with the enzyme activity measurements, Florence Danila and Joanne Lee for TEM, and Hilary Stuart-Williams for the nitrogen elemental analysis. We thank the Centre for Advanced Microscopy at the Australian National University and the Australian Plant Phenomics Facility supported under the National Collaborative Research Infrastructure Strategy of the Australian Government. This research was supported by the Australian Research Council Centre of Excellence for Translational Photosynthesis (CE140100015). CB was funded by H2020 Marie Skłodowska-Curie individual fellowship (DILIPHO, ID: 702755).

## Supplementary information

**Table S1.** Analysis of the decay kinetics of flash-induced variable fluorescence in the absence or presence of 20 μmol DCMU (Fig. 5, left panels) from mesophyll (M) and bundle sheath (BS) thylakoid membranes of S. *viridis* plants grown at high light (HL, 1000 μmol m^−2^ s^−1^) or low light (LL, 300 μmol m^−2^ s^−1^). Nd, not detected.

|  | M LL | BS LL | BS HL | M LL + DCMU | BS LL + DCMU | BS HL + DCMU |
| --- | --- | --- | --- | --- | --- | --- |
| $t_1 (\mu\text{s})$ | 244 | 397 | 433 | nd | <100 | <100 |
| $\text{Ampl}_1 (\%)$ | 73 | 69 | 56 | nd | 26 | 35 |
| $t_2 (\text{ms})$ | 12 | 26 | 50 | 200 | 146 | 127 |
| $\text{Ampl}_2 (\%)$ | 14 | 23 | 22 | 27 | 27 | 38 |
| $t_3 (\text{s})$ | 4.27 | 5.11 | 6.15 | 3.61 | 5.67 | 7.69 |
| $\text{Ampl}_3 (\%)$ | 13 | 21 | 22 | 72 | 47 | 27 |

**Fig. S1.**
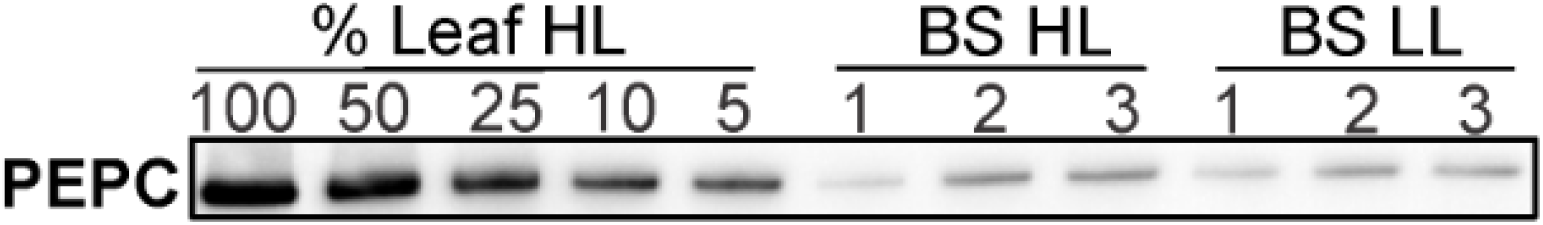
Contamination of bundle sheath (BS) preparations with mesophyll cells assessed by immunoblotting with antibodies against PEP carboxylase (PEPC). HL, *S. viridis* plants grown at high light (1000 μmol m^−2^ s^−1^); LL, plants grown at low light (300 μmol m^−2^ s^−1^). Three biological replicates of BS preparations analysed for each light regime. Samples were normalised on chlorophyll (*a+b*) basis and a dilution series of HL leaf protein sample was used for relative quantification.

**Fig. S2.**
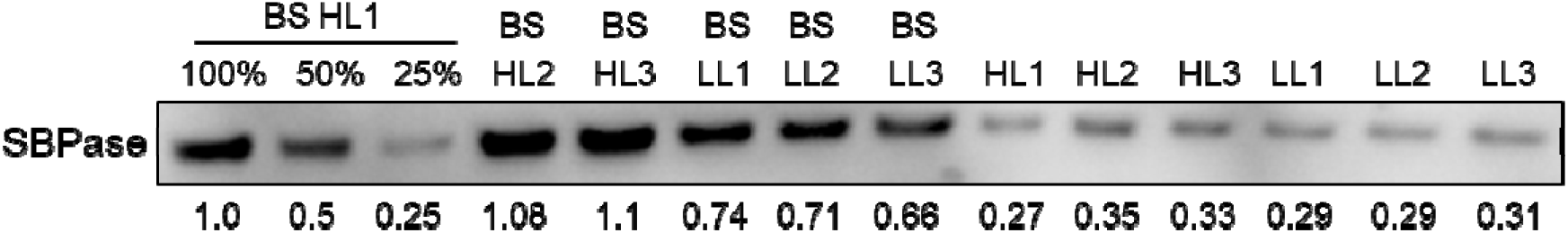
Immunodetection of SBPase in protein samples normalised on chlorophyll (*a+b*) basis. BS HL1-3, bundle sheaths of HL plants; BS LL1-3, bundle sheaths of LL plants; HL1-3 and LL1-3, leaf samples. Quantified relative abundances of SBPase in all samples are shown underneath the blot and the calculated BS chlorophyll values are shown in Table 1.

**Fig. S3.**
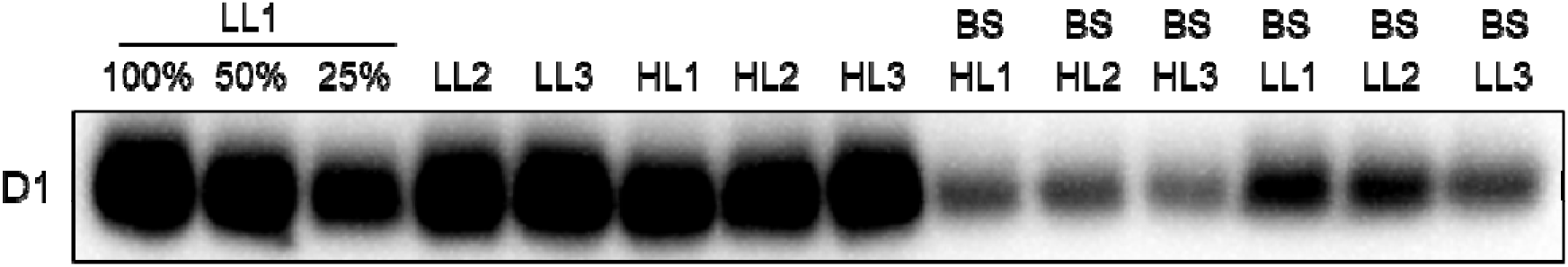
Immunodetection of D1 in bundle sheath (BS) and leaf protein samples normalised on chlorophyll (*a+b*) basis. BS HL1-3, bundle sheaths of HL plants; BS LL1-3, bundle sheaths of LL plants; HL1-3 and LL1-3, whole leaf samples.

**Fig. S4.**
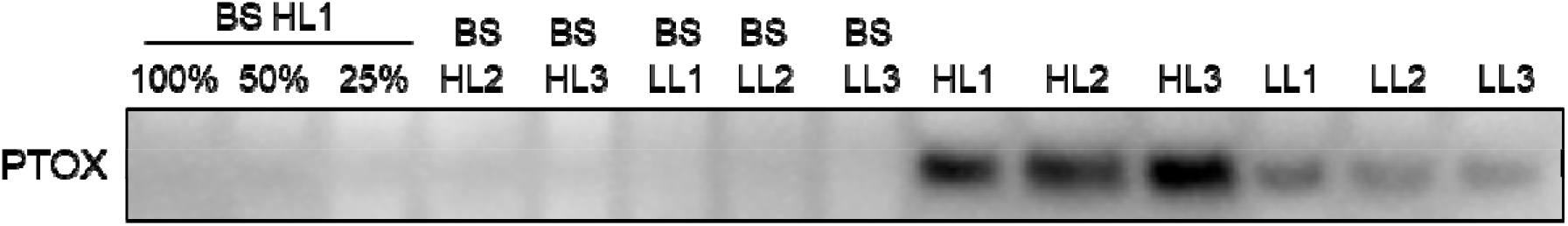
Immunodetection of plastid terminal oxidase (PTOX) in bundle sheath (BS) and leaf protein samples normalised on chlorophyll (*a+b*) basis. BS HL1-3, bundle sheaths of HL plants; BS LL1-3, bundle sheaths of LL plants; HL1-3 and LL1-3, whole leaf samples. No PTOX was detected in BS cells.

